# Updated Database and Evolutionary Dynamics of U12-Type Introns

**DOI:** 10.1101/620658

**Authors:** Devlin C. Moyer, Graham E. Larue, Courtney E. Hershberger, Scott W. Roy, Richard A. Padgett

**Affiliations:** Department of Cardiovascular and Metabolic Sciences, Lerner Research Institute, Cleveland Clinic Lerner College of Medicine, Cleveland Clinic and Department of Molecular Medicine, Case Western Reserve University, Cleveland, Ohio 44106, USA; Department of Molecular and Cell Biology, University of California, Merced, California 95343, USA; Department of Biology, San Francisco State University, San Francisco, California 94132, USA

## Abstract

During nuclear maturation of most eukaryotic pre-messenger RNAs and long non-coding RNAs, introns are removed through the process of RNA splicing. Different classes of introns are excised by the U2-type or the U12-type spliceosomes, large complexes of small nuclear ribonucleoprotein particles and associated proteins. We created intronIC, a program for assigning intron class to all introns in a given genome, and used it on 24 eukaryotic genomes to create the Intron Annotation and Orthology Database (IAOD). We then used the data in the IAOD to revisit several hypotheses concerning the evolution of the two classes of spliceosomal introns, finding support for the class conversion model explaining the low abundance of U12-type introns in modern genomes.

## INTRODUCTION

The process of RNA splicing is a necessary step in the maturation of nearly all eukaryotic pre-messenger RNAs and many long non-coding RNAs. During this process, introns are excised from primary RNA transcripts, and the flanking exonic sequences are joined together to form functional, mature messenger RNAs (1, 2). In most organisms, introns can be excised through two distinct pathways: by the major (greater than 99% of introns in most organisms) or minor (less than 1% in most organisms, with some organisms lacking minor class introns altogether) spliceosome. Despite the existence of eukaryotic species lacking the minor spliceosome, many reconstructions have shown that all eukaryotes descended from ancestors that contained minor class introns in their genomes, all the way back to the last eukaryotic common ancestor (3, 4). The minor class introns have consensus splice site and branch point sequences distinct from the major class introns (5, 6). It was originally thought that the two classes of introns were distinguished by their terminal dinucleotides, with introns recognized by the major spliceosome beginning with GT and ending with AG, and introns recognized by the minor spliceosome beginning with AT and ending with AC. However, it was later shown that introns in both classes can have either sets of terminal dinucleotides and that longer sequence motifs recognized by the snRNA components unique to each spliceosome distinguish the two classes of introns, hence the designations of “U2-type” for the major and “U12-type” for the minor spliceosomes (7).

The large-scale and well-organized online databases of genomic data, like Ensembl (8), UCSC (9), and RefSeq (10), do not provide extensive annotation information of intronic sequence in particular. Many databases focusing primarily on intron annotation information were created in the early 2000s, but most are no longer accessible (11–16), and the ones that remain accessible have not been updated in many years (17, 18). The Exon-Intron Database (EID) (14) was one of the most comprehensive and robust databases in this group, and served as a basis for many further investigations into the peculiarities of introns (19–21), including other, more niche intron annotation databases (15, 22). EID was maintained for at least six years, as it was updated in 2006 (23), but it is no longer accessible. Some more recent databases have been created, like ERISdb (24) and JuncDB (25), but they are narrow in scope: ERISdb only annotates splice sites in a selection of plant genomes, and JuncDB annotates splice sites in a wide variety of genomes, but does not have any other easily-accessible intron annotation information. Of all of the databases mentioned above, U12DB (18) and ERISdb alone annotate intron class. Since U12DB has very old annotation data and ERISdb exclusively annotates introns in plant genomes, there is presently no publically available source of current U12-dependent intron annotation for an evolutionarily diverse array of organisms.

Many features of eukaryotic introns have been examined for clues about their evolutionary history. Introns can be assigned to one of three phases based on their position relative to the codons of the flanking exonic sequence: phase 0 introns fall directly between two codons, phase 1 introns fall between the first and second nucleotides of a single codon, and phase 2 introns fall between the second and third nucleotides of a single codon. It has long been noted that introns are not evenly distributed between the three phases (26, 27). In conjunction with sequence biases on the exonic sides of splice sites, the phase biases were frequently cited by both sides of the debate between the proponents of the “exon theory of genes” (the idea that primordial genes arose through exon shuffling and introns originally came into existence to facilitate this) (28) and those who argued that spliceosomal introns are descended from group II introns that invaded the ancestral eukaryotic genome, preferentially inserting themselves into so-called “proto-splice sites” (29–31). Shortly after the discovery of U12-type introns (5), it was noted that the distribution of U12-type introns in the human genome was nonrandom, further complicating the debate around models explaining the origins of introns by requiring them to explain the presence of two classes of introns, the large discrepancy in the numbers of introns in each class, and the nonrandom distribution of U12-type introns (32). Furthermore, the phase biases in U12-type introns were noted to be different from the previously-documented phase biases in U2-type introns (33, 34).

In an effort to address some of the many open questions about intron evolution, we created the Intron Annotation and Orthology Database (IAOD), a database of intron information for all annotated introns in 24 genomes, including plant, fungal, mammalian, and insect genomes. It also uniquely annotates orthologous introns and intron class. The website is publically accessible at introndb.lerner.ccf.org.

## MATERIALS AND METHODS

### Annotating intron class with intronIC

To begin, intronIC identifies all intron sequences in an annotation file by interpolating between coding features (CDS or exon) within the longest isoform of each annotated gene. For each intron, sequences corresponding to the 5’ splice site (5’SS, from -3 to +9 relative to the first base of the intron) and branch point sequence (BPS) region (from -55 to -5 relative to the last base of the intron) are scored using a set of position weight matrices (PWMs) representing canonical sequence motifs for both U2-type and U12-type human introns. A small “pseudo-count” frequency value of 0.001 is added to all matrix positions to avoid zero division errors while still providing a significant penalty for low-frequency bases. For all scored motifs, the binary logarithm of the U12/U2 score ratio (the log ratio) is calculated, resulting in negative scores for introns with U2-like motifs, and positive scores for introns with U12-like motifs. Because U2 introns are not known to contain an extended BPS motif, the PWM for the U2-type BPS is derived empirically using the best-scoring U12 BPS motifs from all introns in the final dataset whose 5’SS U12 scores are below the 95th percentile. To identify the most likely BPS for each intron, all 12-mer sequences within the BPS region are scored and the one with the highest U12 log ratio score is chosen. This initial scoring procedure follows the same general approach used by a variety of different groups for bioinformatic identification of U12-type introns (4, 32–35).

As originally shown by Burge et al., the 5’SS and BPS scores together are sufficient to produce good binary clustering of introns into putative types, due to strong correspondence between the 5’SS and BPS scores in U12-type introns (4, 32). While this general feature of the data has often been employed in the identification of U12-type introns, a variety of different techniques have been used to define the specific scoring criteria by which an intron is categorized as U2- or U12-type. Here, we have implemented a machine learning method which uses support vector machine (SVM) classifiers (36) to assign intron types, an approach which produces good results across a diverse set of species and provides an easy-to-interpret scoring metric.

Our classification method relies upon two pieces of data: PWMs describing sequence motifs for the different subtypes of U2-type (GT-AG/GC-AG) and U12-type (GT-AG/AT-AC) introns, and sets of high-confidence U2- and U12-type intron sequences with which to train the SVM classifier (Fig 1A). Due to the scarcity of bona fide, experimentally-verified U2- and U12-type introns, a certain amount of curation was required to compile type-specific classifier training and scoring data. For the U12-type set, introns from six previously-published studies (18, 37–41) as well as highly-conserved introns from a number of multi-species ortholog alignments were scored using SpliceRack (34) PWMs, and those with 5’SS scores >0 (i.e. 5’SS motifs more similar to U12-type than U2-type) present in at least three different sources were kept for use as U12-type training data. Combining these introns with branch point data from (40), we identified likely U12-type BPS motifs which were then used to generate BPS PWMs, requiring an A at either position +9 or +10 (following Sheth et al. 2006). For the U2-type set, we first collected intron sequences from the yeast *Schizosaccharomyces pombe*, a species which is believed to lack U12-type introns. These introns were then filtered using data from (42) to include only those with direct evidence of splicing, and scored against human SpliceRack PWMs to establish an upper bound for SpliceRack U12 PWM scores on high-confidence U2-type introns. Finally, using a set of introns conserved between human, zebrafish and horseshoe crab we identified human introns found in orthologous groups where every constituent intron had a 5’SS SpliceRack PWM score less than the *S. pombe* U2-type threshold. These human U2-type introns were combined with the U12-type set to build an updated collection of PWMs, and to define positive (U12-type) and negative (U2-type) training sequences for the SVM (Fig 1B).

**Figure 1.**
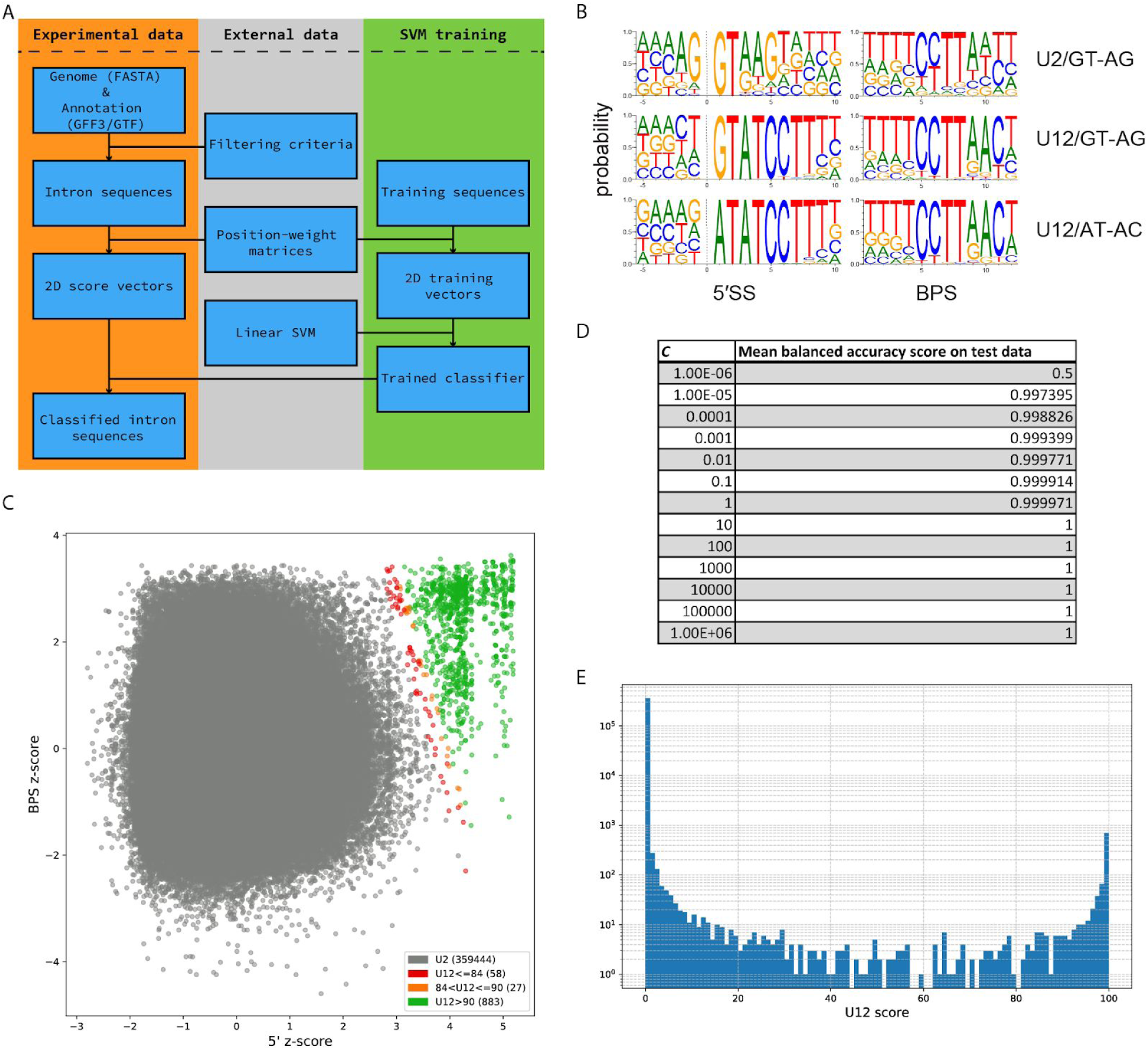
A) Overview of the major steps of the intronIC algorithm B) Sequence logos of the 5’SS and BPS PWMs for GT-AG/AT-AC U2- and U12-type introns C) Scatter plot of all classified introns in the human genome; gray: U2-type introns, red: U12-type introns with probability scores <= 84%; yellow: U12-type introns with probability scores from 84-90%, green: U12-type introns with probability scores > 90%, our chosen scoring threshold D) Balanced accuracy performance of the classifer with different values of hyperparameter *C* on test sets during the first round of the cross-validation process E) Histogram (with logarithmic scale y-axis) of probability scores for the human data shown in part C.

Because clear discrimination between U12-type and U2-type introns can be achieved by considering only two scoring dimensions (4, 32, 34), we use a relatively simple SVM classifier with a linear kernel as implemented in the scikit-learn Python library (43). The SVM is trained on a set of two-dimensional vectors, corresponding to the 5′SS and BPS scores of the introns in the training data, which are labeled by intron type. For linear classifiers there is only a single free hyperparameter to be adjusted, *C*, which is (roughly) the degree to which misclassification of data in the training set is penalized during the creation of an optimized (i.e. wide) margin separating the positive and negative classes. To our knowledge there is no single, standard approach for establishing the best value of *C*; we chose to optimize *C* using an iterative cross-validation method which starts with a wide range of logarithmically-distributed values and narrows that range based upon the best-performing (highest balanced accuracy score) value of *C* in each validation round. After several such iterations, the mean of the resulting range is taken as the final value of *C* to be used to train the classifier. Balanced accuracy is used as a performance metric due to the highly imbalanced nature of the training data, where the negative class (U2-type) greatly outnumbers the positive class (U12-type). Because the human training data is very well-separated, when applied to intron sequences in the human genome values of *C* ≥10 perform equally well during cross-validation (Fig. 1D). Given the broad range of good parameter values, taking the average of all best-performing values results in a more conservative margin (larger *C*) than taking the default “best” parameter value via the scikit-learn API, which simply returns the first rank-1 parameter value found. For the human genome, this approach results in a classifier which performs perfectly on the training sets, with both F1 and precision-recall AUC scores of 1.0 on held-out training data (examples of final scores for human introns in Figs 1C,E).

For the purpose of populating the IAOD, intronIC was slightly modified to produce a single output file containing all of the annotation information recorded in the IAOD for each intron—the default version is available for download from (https://github.com/glarue/intronIC) and the modified version used for this application is available at (https://github.com/Devlin-Moyer/IAOD).

The annotation and sequence files provided as input to intronIC were downloaded from release 92 of Ensembl (with the exception of the FUGU5 assembly of the *Takifugu rubripes* genome, which was downloaded from release 94), release 39 of Ensembl Metazoa, or release 40 of Ensembl Plants (8). Data was obtained for every genome annotated by U12DB with the addition of *Zea mays, Oryza sativa, Glycine max*, and *Schizosaccharomyces pombe* to increase the evolutionary diversity of the represented genomes.

### Annotating introns in non-coding transcripts / regions

To annotate introns, intronIC can use either exon or CDS entries in a GFF3 or GTF file. When using exon entries to define introns, intron phase is undefined.. In order to get complete annotation of both introns within open reading frames and within untranslated regions or non-coding transcripts, intronIC was run twice on each genome analyzed, once producing exon-defined introns and once producing CDS-defined introns. A custom Python script then compared both lists of introns to produce a single list where the CDS-defined intron annotation information was used if the intron was in a coding region and the exon-defined information was used otherwise.

### Finding gene symbols

The output of intronIC includes the Ensembl gene ID but not the gene symbol for all introns in a genome using an annotation file from Ensembl. Ensembl maintains vast databases of genomic data which are accessible with BioMart (44). BiomaRt (45) is an R package for interacting with these databases. A custom R script submitted a list of all of the Ensembl gene IDs in each genome in the database to biomaRt and obtained gene symbols for all of those genes.

### Assigning orthologous introns

Coding sequences for every annotated transcript in each of the 24 genomes were extracted and translated into their corresponding protein sequences. These sequences were aligned with DIAMOND (46) to identify sets of best reciprocal hits—considered as orthologs going forward—between every pairwise combination of species, using an E-value cutoff of 10^−10^ and --min-orfs set to 1. Every pair of orthologous transcripts was then globally aligned at the protein level using Clustal W (v2.1; ref. (47)), and all introns in regions of good local alignment between pairs (≥40% matching amino acid sequence ±10 residues around each intron) were extracted using custom Python scripts (following the approach of ref. (48)). Lastly, conservative clustering of the pairwise orthologous intron sets was performed through identification of all complete subgraphs where every member is an ortholog of every other member (i.e. maximal clique listing) to produce the final intron groups (e.g. A-B, A-C, B-C, B-D → A-B-C, B-D).

### Database creation

A custom Python script created a PostgreSQL database using the output of intronIC, the lists of gene symbols from BioMart, and the list of orthologous groups of introns. All of the orthologous groups were inserted in a table with two columns: a unique numeric ID for each group and a list of all intron labels belonging to that group. One table for each genome contains, for each intron: the abbreviated sequence (see above), taxonomic and common names of the organism, name of the genome assembly, intronIC score, intron class (determined from the intronIC score), intronIC label, chromosome, start coordinate, stop coordinate, length, strand, rank in transcript, phase, terminal dinucleotides, upstream exonic sequence (50 nt), 3’ terminus with the branch point region enclosed with brackets (40 nt), downstream exonic sequence (50 nt), full intron sequence, Ensembl gene ID, Ensembl transcript ID, and gene symbol. Another table with identical fields contains all U12-type introns from all genomes.

### Website design

The website was constructed using Django 2.0, an open-source Python web development framework, and Bootstrap 4.0.0, an open-source framework for front-end web development. The search engines use the Django ORM to interact with the PostgreSQL database.

There are four search engines on the website: the main, advanced, U12, and orthologous searches. The main and advanced search interfaces have input fields corresponding to individual columns in the database, so the text input in each field can easily be matched with the appropriate column using the Django ORM. The U12 search engine uses PostgreSQL search vectors to allow users to make full text queries against the database. I.e. users can input a string containing one term corresponding to as many fields as they like and get a result. However, if the search query contains, e.g., the names of two different species or genes, no results will be returned, since no single record (intron) in the database corresponds to multiple species or genes. This limits the number of possible queries, but allows for a simple user interface for simple queries concerning U12-type introns. Since the main and advanced search engines require users to specify which field of the database each term of their query corresponds to, it does not need to use full text search vectors, and can consequently accept multiple search terms for each field. The homolog search engine also makes use of PostgreSQL search vectors to find the row of the homolog table containing the intron ID input by the user.

### Assessing Randomness of Distribution of U12-type Introns

If U12-type introns were randomly inserted a genome, we would expect the distribution of U12-type introns per gene to be binomial with parameters n = number of genes with at least one U12-type intron and p = 1 - (1 - x)^m-1^, where x is the proportion of U12-type introns in the genome and m is the average number of introns in the genome. Data from the IAOD was used to obtain n, x, and m for each genome in the database that contained at least one U12-type intron. The dbinom function in R was used to compute the probability of observing the observed number of genes with multiple U12-type introns in each genome. Table S1 lists the parameters passed to the dbinom function.

To ensure that the observed clustering of U12-type introns in the same genes was not an artefact of U12-type introns with alternative splice sites being recorded as distinct U12-type introns, all intron coordinates listed by intronIC were used to create a graph where each node corresponded to a position within each genome (e.g. GRCh38+chr1+492045 corresponds to base pair 492045 on chromosome 1 in assembly GRCh38 of the human genome) and two nodes are joined with an edge if they appear in the same row of the list of intron coordinates. In this graph, alternatively spliced introns are evident as clusters of more than 2 nodes, so each cluster represents a single intron, regardless of how many alternative splice sites it possesses. A single edge from each cluster was selected and the corresponding coordinates were matched to the original intronIC output to get accurate counts of the total number of unique introns in each class in all genomes annotated in the IAOD.

## RESULTS AND DISCUSSION

We used intronIC to perform genome-wide identification of U2- and U12-type spliceosomal introns in 24 eukaryotic species including 14 vertebrate animals, 5 invertebrate animals, 4 plants and two yeasts. High-confidence U2- and U12-type introns in human were curated from multiple sources and used to generate type-specific position-weight matrices (PWMs) for the 5’ splice site and branch point sequences. These PWMs were used to create score vectors for every intron in each genome, which were then compared against the high-confidence sets using a machine-learning classifier to assign each intron a probability of being U12-type. Once trained on the conserved intron data, the SVM classifier assigned every intron in the experimental set a probability of being U12-type. Introns with at least a 90% probability of being U12-type are classified as U12-type, which produces classifications in good agreement with previously-reported findings. For example, running our method on U12-type intron sequences from the U12DB (18) results in equivalent classifications for 96% (381/398) of the U12DB introns in chicken, 97% (535/554) in mouse, 94% (15/16) in *Drosophila melanogaster* and 95% (656/691) in human. In *Arabidopsis thaliana*, our method matches the calls in the U12DB 94% of the time (223/238), with similar results (269/292, 92%) for U12-type introns from the plant-specific database ERISdb (24). Furthermore, in each test species listed above intronIC identifies additional putative U12-type introns not present in existing databases, likely due to a combination of newer annotation data and our method’s sensitivity. In *Caenorhabditis elegans*, a well-annotated species believed to have lost all of its U12-type introns, when run on all introns (not just those from the longest isoform per gene) our method categorized only 1/116241 introns as U12-type, suggesting a false-positive rate of less than 0.001%. A total of 8,967 U12-type introns were identified using this technique. Groups of analyzed introns in conserved regions of homologous genes were also annotated; collectively, these data constitute the Intron Annotation and Orthology Database (IAOD).

Figure 2 compares the number of U12-type introns annotated in each species in the IAOD with the numbers of U12-type introns annotated by previous databases annotating U12-type introns: U12DB (18), SpliceRack (34), and ERISdb (24). The IAOD often annotates many more introns than U12DB, likely due to the different approaches to annotating intron class and the quality of the genome assemblies used. In U12DB, U12-type introns were annotated by mapping a set of reference introns from *Homo sapiens, Drosophila melanogaster, Arabidopsis thaliana*, and *Ciona intestinalis* to the whole genomes of every other organism in the database (18), while introns in the IAOD were annotated directly from every genome in the database using the intron-classifying program intronIC (see Methods for details). U12DB primarily annotates U12-type introns in all represented species that are orthologous to the reference U12-type introns (18), while the IAOD annotates U12-type introns in all genomes independently, using the species-specific annotations for each genome. Furthermore, the genome assemblies and annotations used to identify introns in the present study are all several versions newer than those used in U12DB, so part of the discrepancy in the number of U12-type introns annotated is likely due to an increase in the number of annotated genes and splice sites since the creation of U12DB. While intronIC does not provide homology information about the annotated introns, the IAOD also annotates intron orthologs: of the 3,645,636 total introns in the IAOD, 54% (1,989,840) have at least one other intron annotated as being in a conserved region of a homologous gene in another genome in the IAOD.

**Figure 2.**
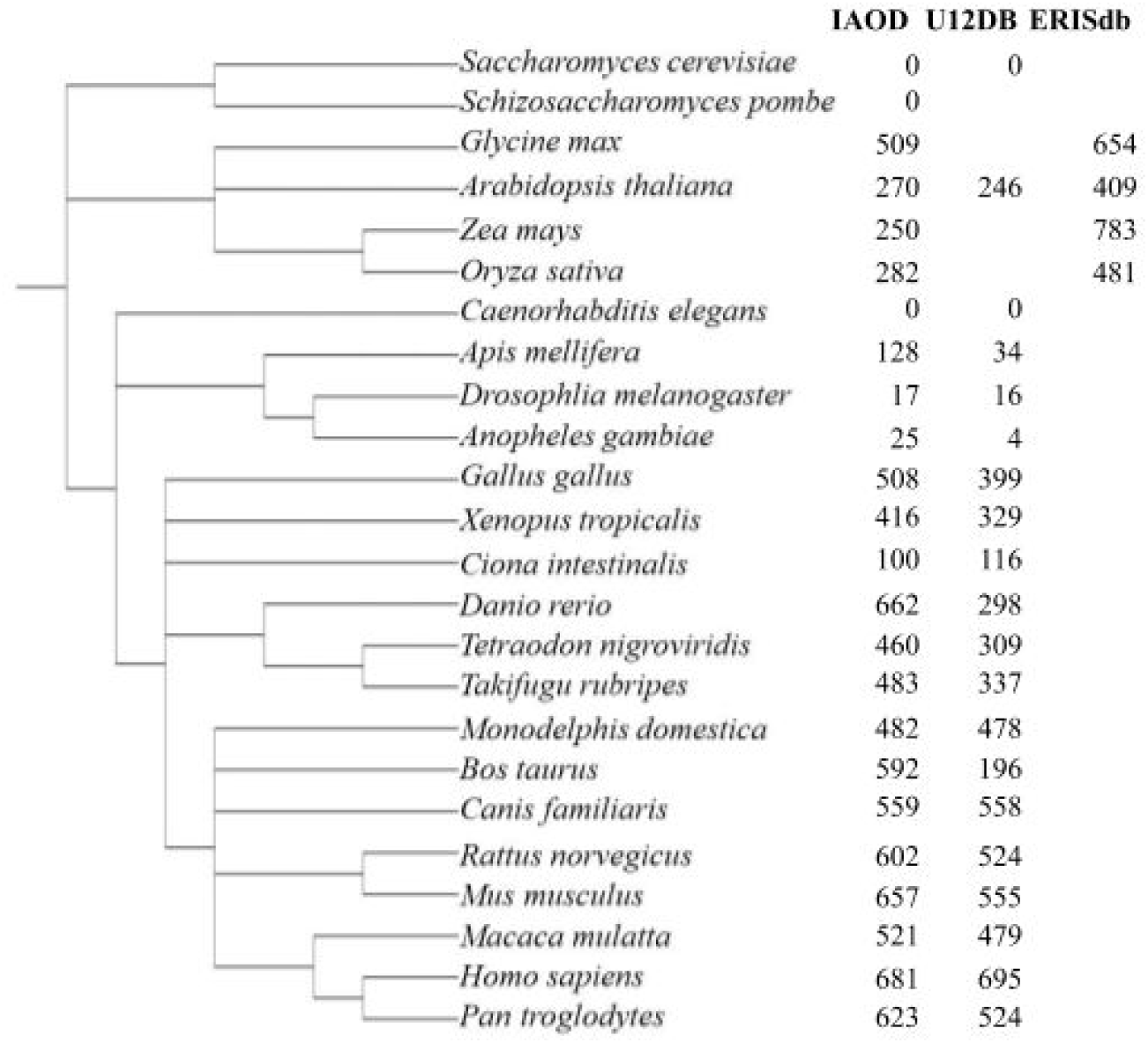
Phylogenetic distribution of U12-type introns in all species annotated by the Intron Annotation and Orthology Database (IAOD), U12DB (18), SpliceRack (34) and ERISdb (24). Blank entries in the table represent organisms not represented in the respective database. Counts of U12-type introns in the IAOD only represent introns flanked by coding exons. The NCBI Taxonomy Browser (69) and Integrative Tree of Life (70) were used to create the phylogenetic tree.

As shown in Figure 2, there are substantially fewer U12-type introns in the analyzed invertebrate animals than in the vertebrates, and none in either species of yeast analyzed, consistent with earlier findings (32, 35). The numbers of introns determined by intronIC to be U12-type in a few species deserve special attention. The numbers of U12-type introns in *A. thaliana, O. sativa* and *Z. mays* are noteworthy because there are substantially fewer U12-type splice sites annotated in the IAOD than in ERISdb, but inspection of the U12-type splice sites annotated in ERISdb reveals many duplicate sequences. These duplicates arise from the fact that ERISdb counts each set of U12-type splice sites from every transcript of every gene as a distinct set of U12-type splice sites. In the case of *A. thaliana*, of the 414 U12-type splice sites annotated in ERISdb, there are only 292 unique sequences, which is much closer to the 269 annotated by the IAOD.

The phase biases observed in the IAOD (Figure 3) agree with the results of previous studies and extend them to many more organisms: an excess of phase 0 introns among U2-type introns (26, 27, 32, 33, 49), and a bias against phase 0 introns among U12-type introns (32, 34) are seen in all studied lineages, and the presence of these biases in both plant and animal genomes suggest a deep evolutionary source. Multiple explanations for the overrepresentation of phase 0 U2-type introns have been proposed, including exon shuffling (50), insertion of introns into proto-splice sites (29, 49), and preferential loss of phase 1 and 2 introns (6). These models do not consider or explain the underrepresentation of phase 0 U12-type introns.

**Figure 3.**
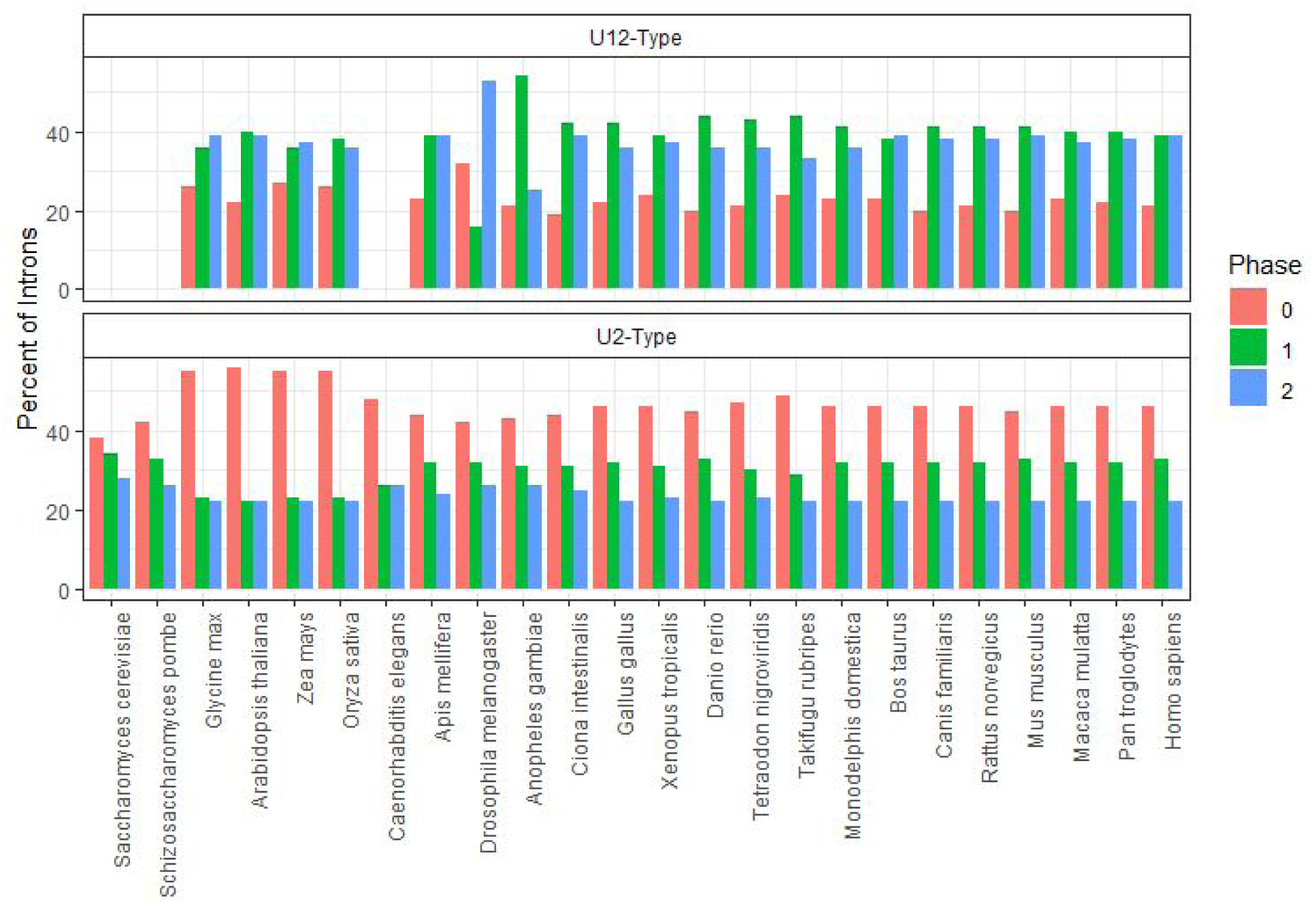
Phase distribution of introns within each class in all genomes annotated in the IAOD. Organisms are grouped by phylogeny. The bias against phase 0 U12-type introns is statistically significant in all organisms but *G. max, O. sativa, X. tropicalis*, and *Z. mays* (chi-squared; p < 0.10). The bias towards phase 0 U2-type introns is statistically significant in all organisms but *A. mellifera, D. melanogaster, S. cerevisiae*, and *S. pombe* (chi-squared; p < 0.10).

To account for the phase biases present in U12-type introns, we propose that the observed phase biases in both classes of introns can be explained by an extension of the class-conversion hypothesis proposed by Burge et al. (32). This hypothesis arose from the observation that U12-type introns in human genes were often found to have U2-type introns at orthologous positions in *C. elegans* genes. Dietrich et al. (7) showed that U12-type introns could be converted to U2-type introns with as few as two point mutations. These results also suggest that class conversion is likely to only proceed from U12-type to U2-type. In light of this, one possible explanation for the current data is that, at an early stage in eukaryotic evolution, there were many more U12-type introns than are currently observed in any characterized genome, and the phase bias arose as phase 0 U12-type introns were preferentially converted into U2-type introns, producing both an overrepresentation of phase 0 U2-type introns and an underrepresentation of phase 0 U12-type introns. This selectivity for phase 0 introns in the class conversion process rests on the function of the -1 nucleotide relative to the 5’ splice site in both spliceosomes.

As shown in Figure 4, there is a large excess of G at the -1 position of U2-type 5’ splice sites, across all three phases, in agreement with earlier investigations (50, 51). Figure 4 also shows that there is an excess of -1U in U12-type introns in all three phases. There also appears to be a bias against -1 A and G in phase 1 and phase 2 U12-type introns, but not in phase 0 U12-type introns. Interestingly, when introns are grouped by terminal dinucleotides, these biases are only found in U12-type introns with GT-AG terminal dinucleotides and not in U12-type AT-AC introns (Figure 4). The preference for -1G at U2-type 5’ splice sites appears to be due to the fact that the -1 nucleotide pairs with a C on the U1 snRNA (52, 53). The preference for -1U at U12-type 5’ splice sites is more mysterious, as no snRNAs are known to bind to this position. It was previously shown that the U11/U12-48K protein interacts with the +1, +2 and +3 nucleotides at the U12-type 5’ splice site in a sequence-specific fashion, but the specificity of the interaction with the -1 position was not studied (54). As noted above, the bias against -1G in U12-type introns with GT-AG terminal dinucleotides could be a consequence of the gradual conversion of many U12-type introns into U2-type introns. The lack of consistent -1 nucleotide biases in AT-AC introns of either class may be due to the fact that AT-AC introns are poorly recognized by the U2-type spliceosome (53) and were thus largely unaffected by the class conversion process.

**Figure 4.**
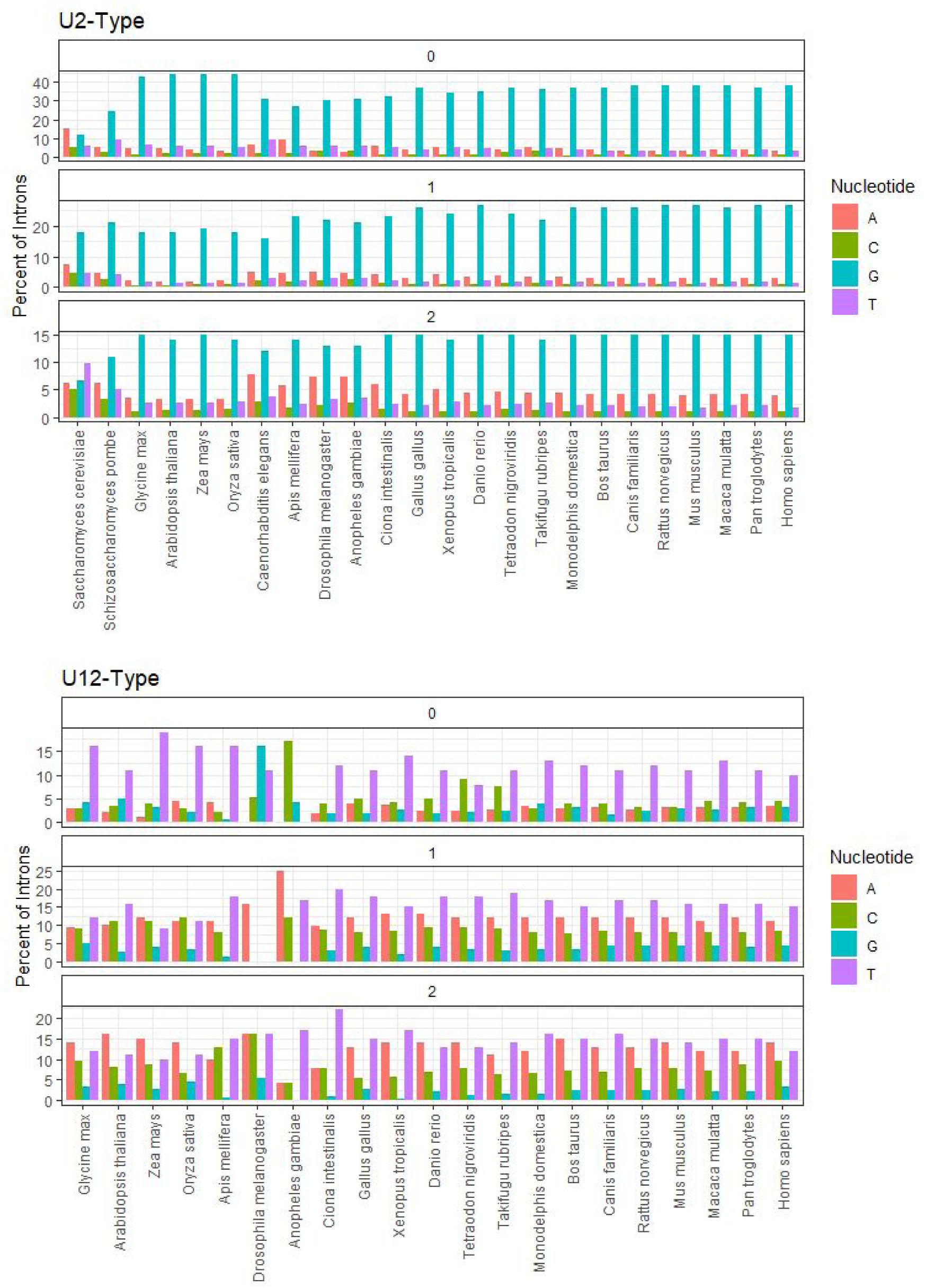
Percentages of introns with the specified nucleotide immediately upstream of the 5’ splice site in each phase of both classes of introns. Organisms are grouped by phylogeny.

We propose that this preference for conversion of phase 0 U12-type introns is due to the fact that introns with a G at the -1 position relative to the 5’ splice site bind more strongly to the U1 snRNA (53, 54), and the -1 nucleotide of phase 0 introns is the final wobble position of the corresponding codon and can be a G in 13 of 20 codon families. Thus, the -1 nucleotides of phase 0 U12-type introns were more free to mutate to G and increase the affinity of the U2-type spliceosome for their 5’ splice sites, gradually accumulating mutations in the other sequences required for recognition by the U12-type spliceosome (7). Table 1 contains some examples of orthologous introns of different classes that demonstrate the class conversion process. This unidirectional conversion process also provides an explanation for the low abundance of U12-type introns in modern eukaryotic genomes.

**Table 1.**
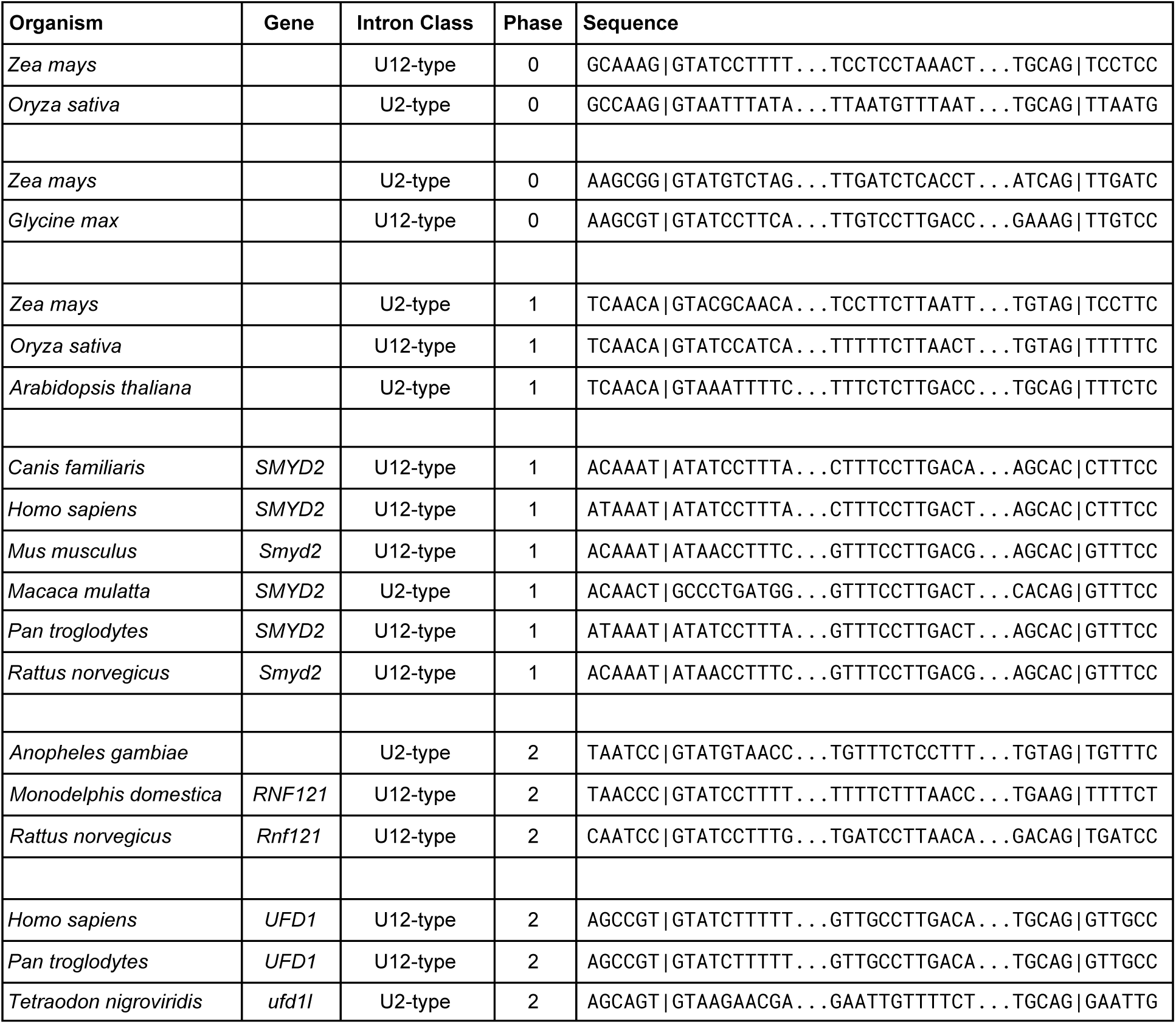
Groups of orthologous introns of different classes. Introns with nothing in the gene column came from transcripts with no gene name annotated by Ensembl. Vertical bars in the sequence column denote splice sites, and the middle sequence flanked by ellipsess is the putative branch point region as annotated by intronIC.

Multiple previous surveys of U12-type introns have revealed that the distribution of U12-type introns in the human genome is non-random, i.e. there is a statistically significant tendency for U12-type introns to cluster together in the same genes (32–34). Repeating this analysis for all genomes in the IAOD (except *S. cerevisiae, S. pombe*, and *C. elegans*, as they lack U12-type introns) replicated their findings in all 21 genomes (p < 0.05 for all genomes; see Methods and Table S1). Many explanations for this nonrandom distribution have been proposed, including the fission-fusion model of intron evolution (32); a difference in the speed of splicing of U12-type and U2-type introns (34, 38); and the idea that the U12-type introns arose during an invasion of group II introns after U2-type introns had already seeded the ancestral eukaryotic genome, meaning the new U12-type introns could only be inserted in certain locations (55). The fission-fusion model posits that two separate lineages of the proto-eukaryote evolved distinct spliceosomes and then fused their genomes such that all genes originally contained either only U2-type introns or only U12-type introns. Thus, modern U2-type introns in genes also containing U12-type introns were originally U12-type introns that were subjected to the class conversion process discussed above (32). An alternative argument for the low abundance of U12-type introns is that they are excised more slowly than U2-type introns, so genes that contain U12-type introns contain them because those genes need to be expressed slowly for some reason (34, 38). However, it has since been shown that the rate of excision of U12-type introns is not sufficiently different from the rate of excision of U2-type introns to produce a meaningful impact on the expression of transcripts containing U12-type introns (56). Furthermore, recent evidence suggests that the rates of both types of splicing are sufficiently fast that most introns will be excised cotranscriptionally (39).

Basu et al. (57) argued that the number of U12-type introns present in the ancestral eukaryotic genome was unlikely to be substantially larger than the largest number of U12-type introns observed in any modern genome, thus suggesting that the process of class conversion is a minor evolutionary force. However, the basis of their argument is the finding that the positions of U12-type introns are more highly conserved than the positions of U2-type introns between humans and *Arabidopsis thaliana*, a result that the present data do not support: we find that out of the 93 U12-type introns in the human genome in regions of good alignment to *A. thaliana*, only 8 (9%) are in conserved positions, while out of the 9,527 U2-type introns in such regions of the human genome, 2,098 (22%) are in conserved positions in *A. thaliana*. Thus, our comparative analysis is consistent with U12-type intron enrichment in the ancestral eukaryotic genome relative to the most U12-intron-rich extant lineages.

The majority of introns annotated in the IAOD in both classes begin with GT and end with AG (Table 2), in agreement with previous studies (17, 32, 34). A substantial minority of U2-type introns, but almost no U12-type introns, were found to have GC-AG as their terminal dinucleotides in many of the analyzed genomes, reflecting their previously documented role in alternative 5’ splice site selection in U2-type splicing in many organisms (6, 34, 58–60). Several previous studies have found numerous introns with other non-canonical terminal dinucleotides in multiple genomes, sometimes with functional roles in regulation of alternative splicing (17, 34, 58, 61), but intronIC has annotated many thousands of U2-type introns with non-canonical terminal dinucleotides in certain organisms, such as *Gallus gallus* and *Tetraodon nigroviridis* (Table 2). Inspection of these introns reveals that the vast majority of these splice sites are only a few nucleotides away from a conventional U2-type splice site with canonical terminal dinucleotides; these splice sites with non-canonical dinucleotides were likely annotated on the basis of conserved exon boundaries, without regard for the precise placement of the splice sites. The proportion of U12-type introns with non-canonical terminal dinucleotides (Table 2) largely agrees with previous investigations (34, 35, 62).

**Table 2.**
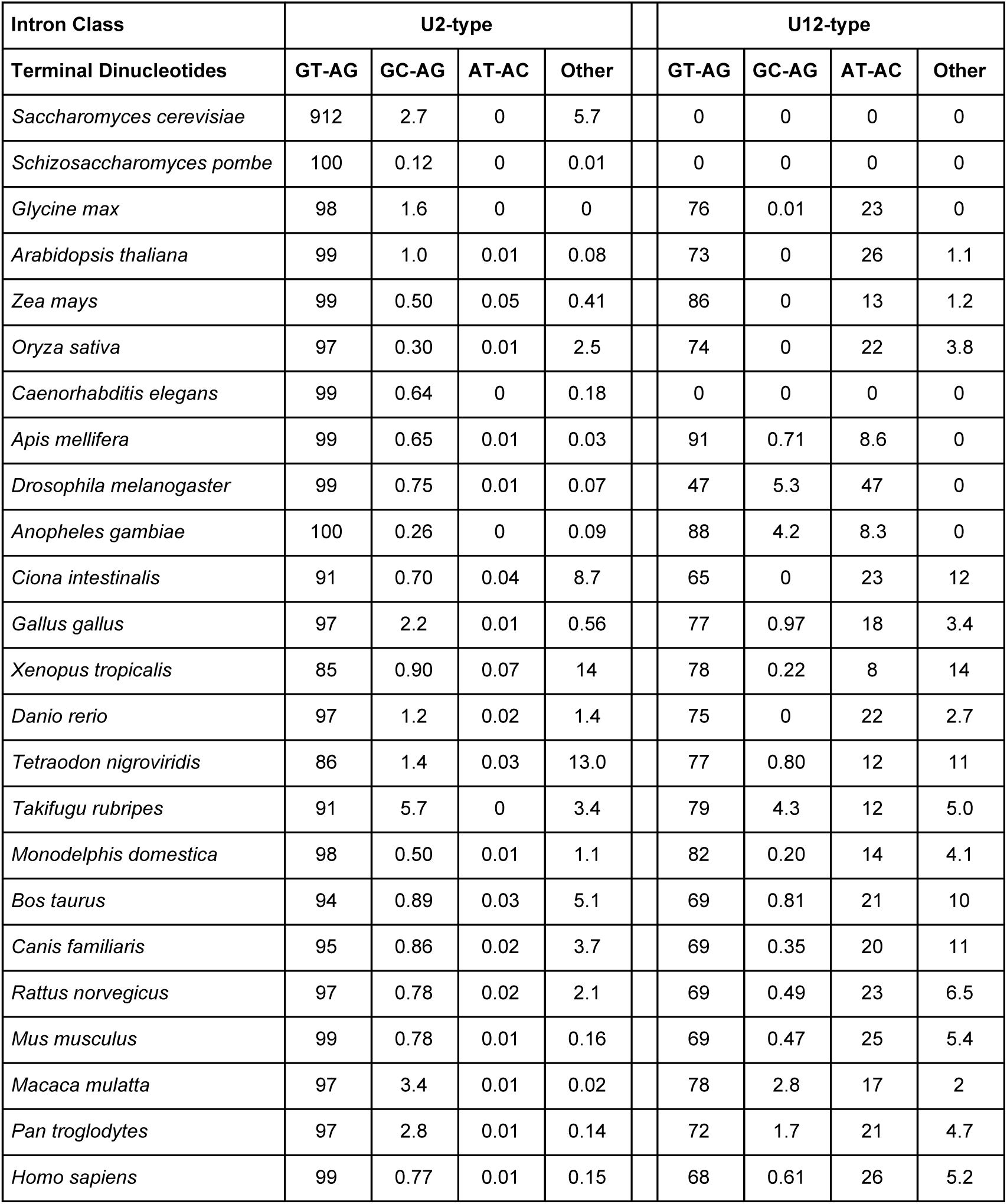
Percentages of introns with various terminal dinucleotides in each class in all annotated genomes. Organisms are sorted in the phylogenetic order shown in Figure 2.

Figure 6 shows the distributions of intron lengths in six of the genomes annotated in the IAOD, representing each general type of length distribution observed in the IAOD. In accordance with previous studies, when plotted on a log scale, there are two distinct peaks in the distribution of intron lengths in U2-type introns in humans and chicken while the distribution of U12-type intron lengths has only one peak (33, 63) (these peaks are not apparent when length is plotted on a linear scale). A previous study considered the distribution of intron lengths amongst several eukaryotic genomes collectively (64), producing a distribution similar to those observed in the human and chicken genome in Figure 6. However, Figure 6 demonstrates great diversity in the distributions of intron lengths amongst eukaryotes; zebrafish have two distinct peaks of comparable size of intron lengths in both classes, while corn, honeybee and fugu have large peaks of shorter introns and very small peaks of longer introns in both classes. The significance of these variations is unclear; differing distributions of intron lengths in the two classes of introns have previously been used to argue that U12-type introns are recognized through intron definition, while U2-type introns are recognized by exon definition (65). However, Figures 7 and 8 show that the mean intron lengths in both classes of intron in all 24 genomes annotated in the IAOD correlate strongly with genome size (Pearson’s r: 0.87 for U12-type introns and 0.93 for U2-type introns), consistent with previous findings (63, 64, 66). This correlation suggests that mean intron lengths in both classes are generally a function of genome size and not a reflection of intron definition imposing a restriction on the size of U12-type introns.

**Figure 5.**
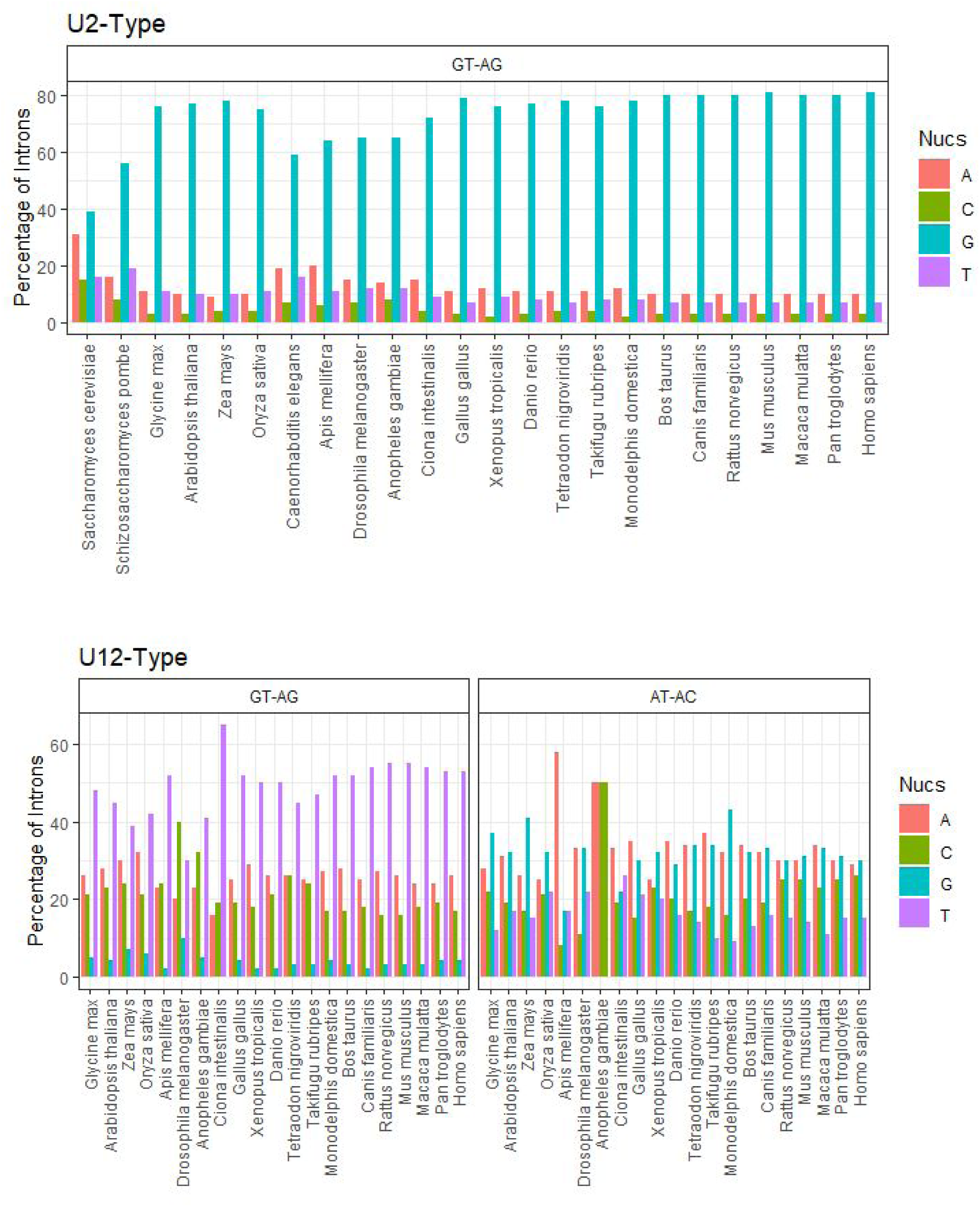
Nucleotide biases at the -1 position relative to the 5’ splice site for all organisms (excluding those lacking U12-type introns) annotated in the IAOD, grouped by terminal dinucleotides and intron class.

**Figure 6.**
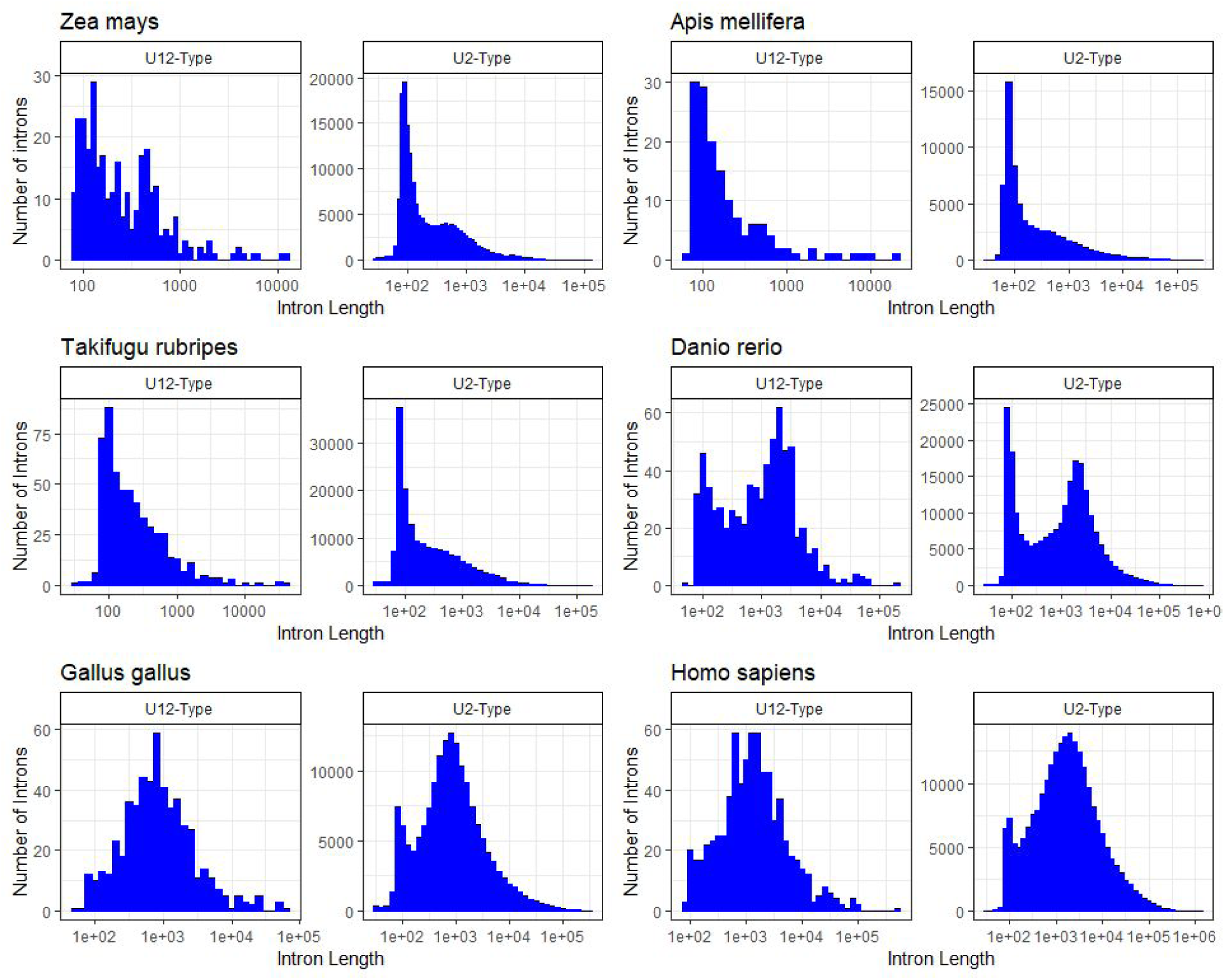
Distributions of intron lengths in both classes of intron in six of the genomes annotated in the IAOD. The x-axis of each plot is a log scale.

**Figure 7.**
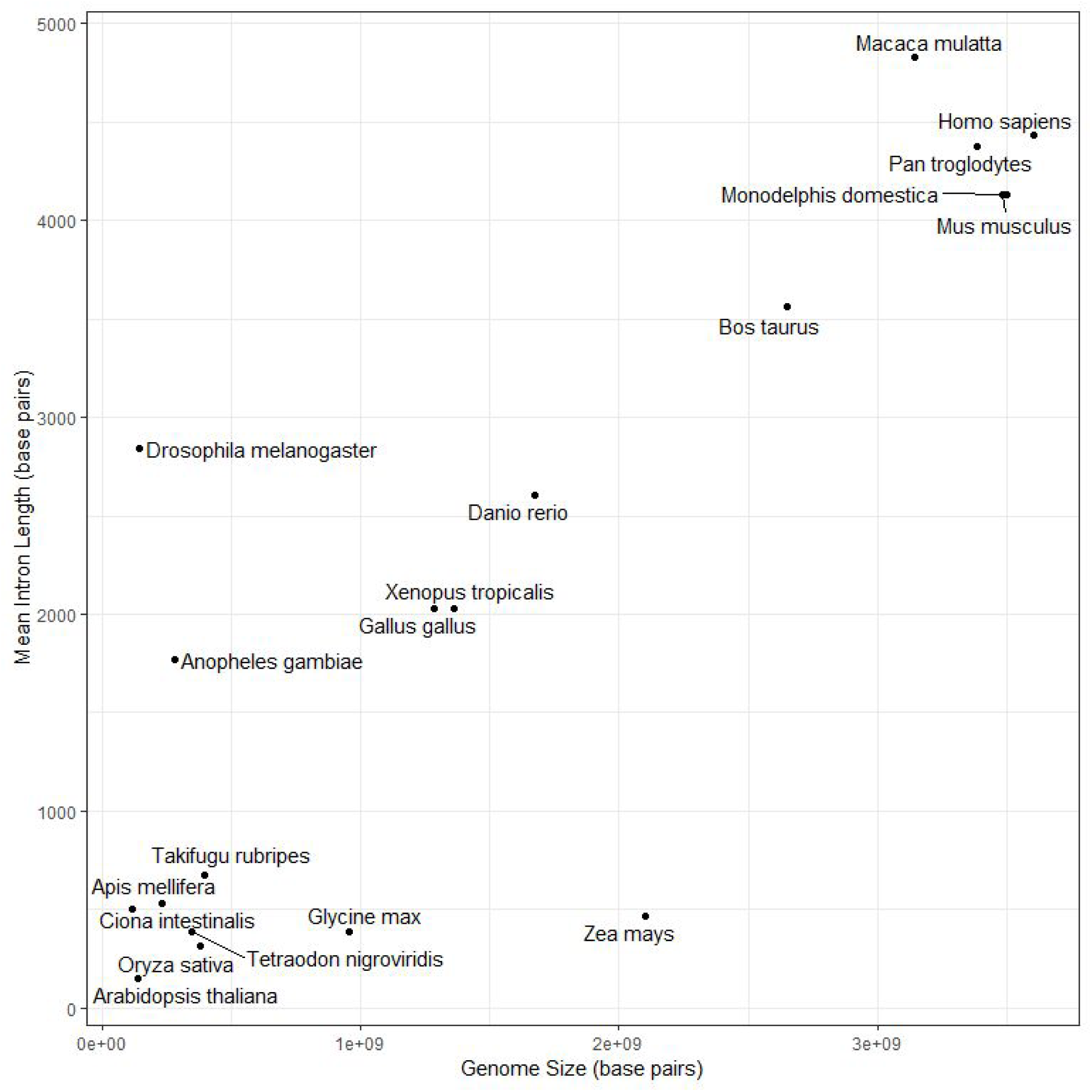
Relationship of genome size and mean U12-type intron length in genomes annotated in the IAOD. *Schizosaccharomyces pombe, Saccharomyces cerevisiae*, and *Caenorhabditis elegans* are not shown in this figure as they lack U12-type introns.

**Figure 8.**
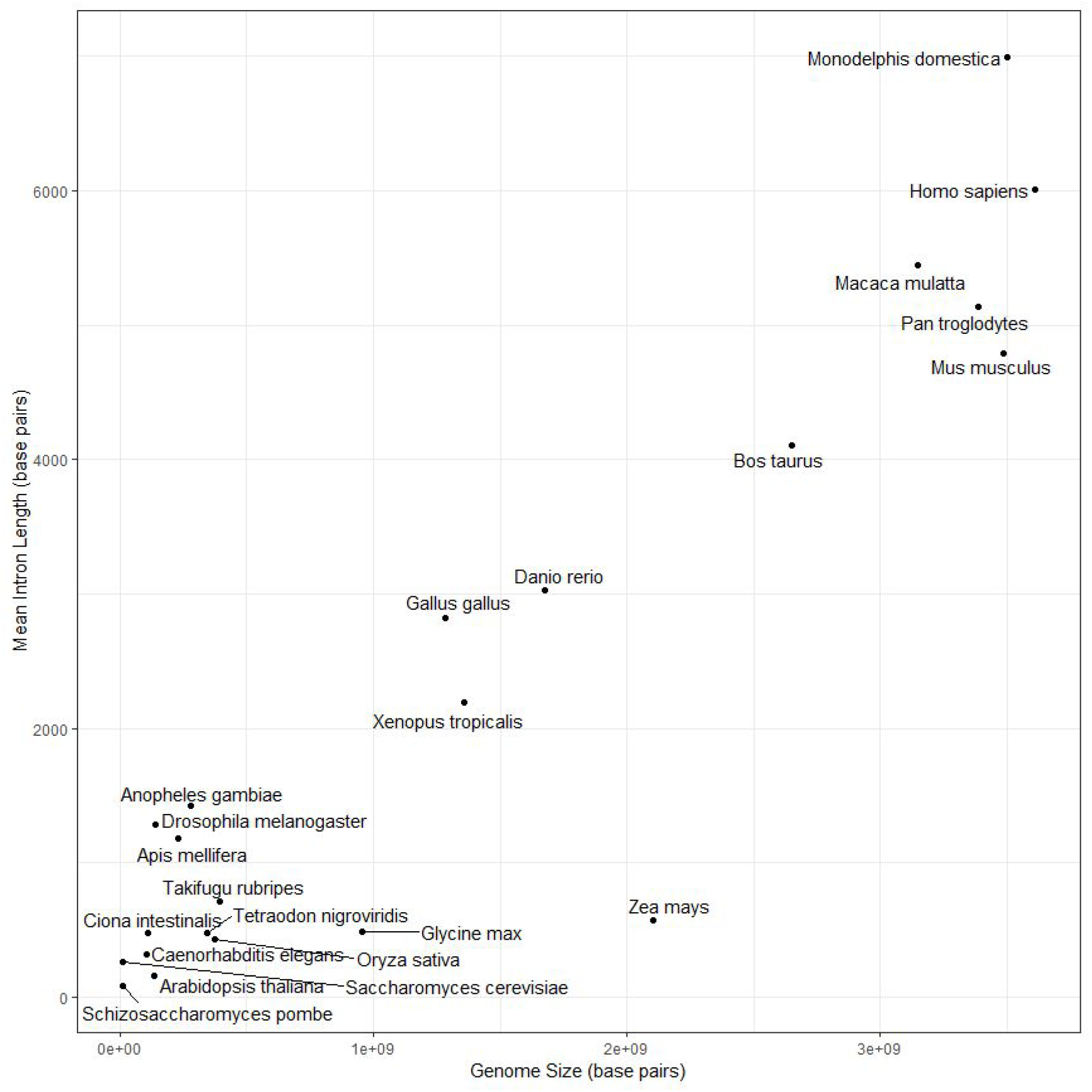
Relationship of genome size and mean U2-type intron length in genomes annotated in the IAOD.

Interestingly, Figures 7 and 8 show that the relationship between mean intron length and genome size differs between vertebrates, insects, and plants. In insects, the total genome size remains very small and the mean intron length does not appear to correlate with total genome size. This may be related to the greater prevalence of intron definition in splicing in insects than in vertebrates (67, 68). In plants, mean intron length does appear to correlate with total genome size, but mean intron length increases much more slowly with total genome size than in vertebrates. Similar correlations are observed between mean intron length and gene number, with a much more prominent difference between the slope of the correlation in plants and vertebrates (data not shown). The significance of this remains unclear.

## CONCLUDING REMARKS

We have created a database of intron annotation and homology information and used it to investigate several evolutionary hypotheses regarding the two classes of spliceosomal introns in eukaryotes. We have also created a web-based interface for querying this database to facilitate further investigations. The relationships between intron class, phase, terminal dinucleotides and -1 nucleotides at the 5’ splice site and the nonrandom distribution of U12-type introns annotated in the IAOD do not support many previous models that explain these patterns (6, 29, 49, 50), but do support an extension of the class conversion model proposed in ref. 33.

## DATA AVAILABILITY

The IAOD is publically accessible at introndb.lerner.ccf.org and all code used to create the database and run the website is available at the following GitHub repository https://github.com/Devlin-Moyer/IAOD.

## AUTHOR CONTRIBUTIONS

DM planned, designed, and created the website and all scripts used to generate and process data except intronIC. GL designed and wrote intronIC. DM wrote the manuscript with input from all authors.

## ACKNOWLEDGEMENTS

Michael Weiner provided invaluable technical support by hosting and managing the website. Rosemary Dietrich provided extensive background on splicing and general advice on various aspects of data interpretation. Daniel Blankenburg provided valuable feedback on various aspects of the web design and method of annotating orthologous introns.

