## Supplemental Table S1 for "Updated Database and Evolutionary Dynamics of U12-Type Introns"

| **Organism** | **Number of U12-Type Introns** | **Genes with Multiple U12-Type Introns** | **Proportion of Introns That Are U12-Type** | **Probability That U12-Type Introns Are Randomly Distributed** |
| --- | --- | --- | --- | --- |
| *Glycine max* | 521 | 25 | 0.0025 | 2.3E-10 |
| *Arabidopsis thaliana* | 274 | 16 | 0.0025 | 3.6E-08 |
| *Zea mays* | 282 | 8 | 0.0017 | 0.0011 |
| *Oryza sativa* | 281 | 12 | 0.0024 | 7.5E-06 |
| *Apis mellifera* | 140 | 4 | 0.0021 | 0.026 |
| *Anopheles gambiae* | 24 | 1 | 6.0E-4 | 0.036 |
| *Ciona intestinalis* | 104 | 4 | 0.0011 | 0.0027 |
| *Gallus gallus* | 503 | 33 | 0.0034 | 1.8E-08 |
| *Xenopus tropicalis* | 422 | 85 | 0.0026 | 8.6E-47 |
| *Danio rerio* | 643 | 43 | 0.0030 | 2.9E-10 |
| *Tetraodon nigroviridis* | 474 | 25 | 0.0028 | 1.4E-4 |
| *Takifugu rubripes* | 521 | 23 | 0.0032 | 0.0025 |
| *Monodelphis domestica* | 470 | 28 | 0.0028 | 2.6E-08 |
| *Bos taurus* | 574 | 40 | 0.0035 | 1.3E-06 |
| *Canis familiaris* | 542 | 44 | 0.0035 | 7.7E-09 |
| *Rattus norvegicus* | 583 | 42 | 0.0034 | 7.7E-10 |
| *Mus musculus* | 630 | 49 | 0.0032 | 2.5E-17 |
| *Macaca mulatta* | 550 | 37 | 0.0033 | 9.9E-09 |
| *Pan troglodytes* | 639 | 45 | 0.0036 | 5.7E-09 |
| *Homo sapiens* | 674 | 48 | 0.0033 | 4.8E-16 |

Table S1. Probabilities that U12-type introns were randomly inserted into each genome along with all parameters used to calculate those probabilities. If U12-type introns were randomly inserted a genome, one would expect the distribution of U12-type introns per gene to be binomial with parameters n = number of genes with a U12-type intron and p = 1 - (1 - x)^m-1^, where x is the proportion of U12-type introns in the genome and m is the average number of introns in the genome. Organisms are grouped by phylogeny. *S. cerevisiae*, *S. pombe*, and *C. elegans* were omitted from this analysis as they lack U12-type introns. *D. melanogaster* was omitted from this analysis as there are no genes with multiple U12-type introns in that genome.
